# Frontoparietal activity related to neuropsychological assessment of working memory

**DOI:** 10.1101/2025.01.13.632797

**Authors:** August Jurva, Balbir Singh, Helen Qian, Zhengyang Wang, Monica L. Jacobs, Kaltra Dhima, Dario J. Englot, Shawniqua Williams Roberson, Sarah K. Bick, Christos Constantinidis

## Abstract

Executive functions, including working memory, are typically assessed clinically with neuropsychological instruments. In contrast, computerized tasks are used to test these cognitive functions in laboratory human and animal studies. Little is known of how neural activity captured by laboratory tasks relates to ability measured by clinical instruments and, by extension, clinical diagnoses of pathological conditions. We therefore sought to determine what aspects of neural activity elicited in laboratory tasks are predictive of performance in neuropsychological instruments. We recorded neural activity from intracranial electrodes implanted in human epilepsy patients as they performed laboratory working memory tasks. These patients had completed neuropsychological instruments preoperatively, including the Weschler Adult Intelligent Scale and the Wisconsin Card Sorting test. Our results revealed that increased high-gamma (70-150 Hz) power in the prefrontal and parietal cortex after presentation of visual stimuli to be remembered was indicative of lower performance in the neuropsychological tasks. On the other hand, we observed a positive correlation between high-frequency power amplitude in the delay period of the laboratory tasks and neuropsychological performance. Our results demonstrate how neural activity around task events relates to executive function and may be associated with clinical diagnosis of specific cognitive deficits.

## INTRODUCTION

The ability to regulate behavior voluntarily by allocating neural resources to tasks and stimuli deemed important in the context of current contingencies is known as executive function, and working memory is an important component of it (Friedman and Robbins, 2022). Many psychiatric disorders and neurological conditions are associated with deficits in working memory and executive function that greatly impact patients’ cognitive abilities and everyday life (Menon, 2011; Sha et al., 2019; Snyder et al., 2015). In the laboratory setting, neural activity associated with executive function is often assessed using tasks that evaluate working memory and context updating; response inhibition and interference control; and mental set shifting (Constantinidis and Klingberg, 2016; Munoz and Everling, 2004). Clinically, deficits in cognition are diagnosed using standardized and norm-referenced neuropsychological instruments, such as the Weschler Adult Intelligence Scale (Wechsler, 2008). These assessments typically contain components that measure executive function components, including working memory, processing speed, and cognitive flexibility. A few tasks have found applications in both clinical settings and laboratory studies, such as the Wisconsin Card Sorting test (Buckley et al., 2009; Heaton and Staff, 1993).

The neural basis of executive functions and particularly working memory has been explored in the laboratory with computerized tasks that involve presentation of sensory stimuli, a delay period over which they have to be maintained in memory, followed by a recall test of some sort (Buschman and Miller, 2022). Studies in animal models of working memory have been especially fruitful in uncovering neural correlates of working memory, although controversy remains about its underlying neural mechanisms (Constantinidis et al., 2018; Miller et al., 2018). However, due to species and methodological differences, it is not well understood what aspects of neural activity are most strongly related to underlying executive function.

We were therefore motivated to investigate what aspects of neural activity elicited by laboratory task events best capture overall working memory ability as evaluated by neuropsychological instruments. In particular, we hypothesized that decreased activation of the prefrontal cortex during the laboratory tests would be predictive of impaired performance in neuropsychological tests, as these brain areas have been implicated in generation of elevated neuronal activity during working memory performance (Riley and Constantinidis, 2016). We recorded neural activity from epilepsy patients implanted with intracranial electrodes while they participated in laboratory visual working memory tasks and sought to uncover the neurophysiology characteristics that best predict clinical metrics and could represent biomarkers of executive function impairment.

## RESULTS

### Behavioral Performance in clinical and laboratory tasks

Neural activity and neuropsychological performance were evaluated in a total of 28 participants (10 male, 18 female – Supplementary Table 1). Preoperatively, patients completed outpatient clinical neuropsychological evaluations that included a battery of tests (discussed below). Each participant was then implanted with sEEG leads, with number and anatomic locations of electrodes determined entirely by clinical considerations related to hypotheses about seizure onset zone. After surgery and during their admission to the epilepsy monitoring unit, patients were tested with spatial and shape visual working memory tasks while activity was recorded from intracranial electrodes (Fig.1). We refer to these as “laboratory tasks” hereafter, as they were designed to mimic tasks typically used in human and animal studies of neural activity associated with working memory in laboratory settings. In the spatial task, the participants had to remember the location of the stimulus presented during the cue period and, after a delay period, indicate where the stimulus appeared by dragging a pointer to the location of the stimulus (Fig. 1A). The spatial stimulus set varied the spatial location of a white disk along an (invisible) circle. Two versions of the spatial task were used, one with a 3s and one with a 6s delay period. The shape working memory task involved different polygon shapes, always presented in the center of the screen (Fig. 1B). Two stimuli were presented in sequence with an intervening delay period between them. The subjects needed to determine whether the two stimuli were identical or not and indicate their judgment by selecting a green or red choice target.

**Figure 1.**
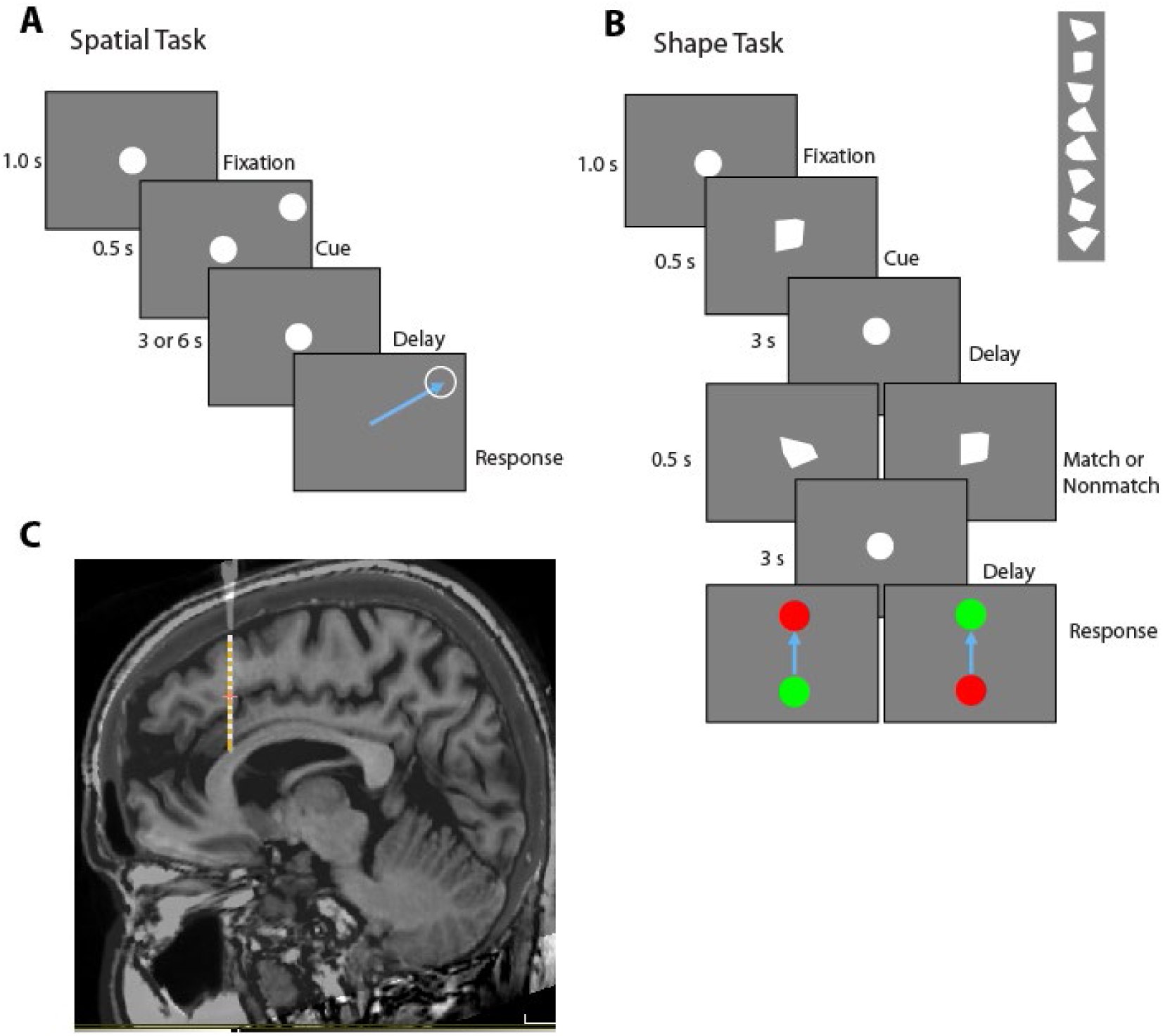
Behavioral tasks and recording methods. A. Spatial Task. At the start of each trial, a white disk appears in the center of the tablet screen, and the subject moves the stylus into it to initiate the trial. After 1 second, a second white disk appears (Cue) at a peripheral location for 0.5 s, after which only the center disk remains. After a delay period of either 3s or 6s, the center disk disappears and the subject needs to drag the stylus across the screen into the remembered location of the cue. B. Shape Task. At the start of each trial, a white disk appears in the center of the tablet screen, and the subject moves the stylus into the circle to initiate the trial. After a delay period, a white polygon replaces the center disk for 0.5 s (Cue), followed by the reappearance of the center disk. After a delay, a second convex polygon replaces the center disk for 0.5 s, followed by the reappearance of the center disk. After a second delay, the center disk disappears and the subject needs to drag the stylus to either a green or red peripheral circle to indicate whether the two polygons were the same or not, respectively. **C.** Example MRI scan of one patient, with electrode position, based on CT scan, superimposed, where LFP recording were made.

In the 3s delay spatial task (n = 17), errors involved incorrect timing (responding too early or too late relative to the offset of the fixation point) or mis-localizing the stimulus. To assess the fidelity of working memory, we examined the mean absolute error distance was 0.95 ± 0.32 cm and precision, defined as the standard deviation of error distance was 0.87 ± 0.63 cm. In the 6s delay spatial task (n = 17) the mean absolute error distance was 0.93 ± 0.66 cm, and the mean precision was 0.87 ± 0.63 cm. The shape task (n = 26) was essentially a two-alternative forced choice task, with responses towards visible choice targets. Working memory fidelity was best captured in this task by the percentage of correct trials in a session, which we refer to as “performance”. The mean performance across subjects was 80.7 ± 10.8%.

Our analysis of performance in the neuropsychological battery focused on tests that assessed executive function ability. These included the Wisconsin Card Sorting Test 64 Card Version (WCST-64) and the Weschler Adult Intelligence Scale Fourth edition (WAIS-IV) mental arithmetic and digit span subtests. The WCST involves matching a target card to one of four category cards based on color, shape, or number. The experimenter changes the classification rule without warning and the participant must identify the new pattern. The digit span task requires subjects to repeat a series of digits of increasing length forward, backward, and sequencing lowest to highest. The mental arithmetic task involves timed arithmetic word problems to be solved mentally without the use of pencil and paper. Raw scores from all tests were converted to z-scores for all comparisons. Mean standardized scores from our sample were -0.45 ± 0.75 (n = 23) for the WCST, -0.56 ± 0.66 (n = 26) for the digit span test, and -0.86 ± 0.68 (n = 26) for the mental arithmetic task.

Performance in the two WAIS-IV tests, mental arithmetic and digit span tests, was positively correlated (r=0.53, p=0.010), whereas the correlation of the WCST task with either of these tests was weaker (r=0.22, p=0.34 with digit span; r=0.07, p=0.075 with mental arithmetic, Fig. 1A). Correlation between neuropsychological and laboratory tests (Fig. 2A) was highest for precision in the 6s spatial memory task and performance in the digit span task (r=0.78, p=2.6e-3). More modest correlation was observed between precision in the 6s spatial working memory task and performance in the mental arithmetic task (r=0.39, p=0.20), and between performance in the digit span and shape working memory tasks (r=0.29, p=0.21). We note that the distribution of patient performance was skewed to below mean population values for all three neuropsychological tests (Fig. 2B), as can be expected for this patient population given the high rates of cognitive dysfunction with medically refractory epilepsy (Stretton and Thompson, 2012).

**Figure 2.**
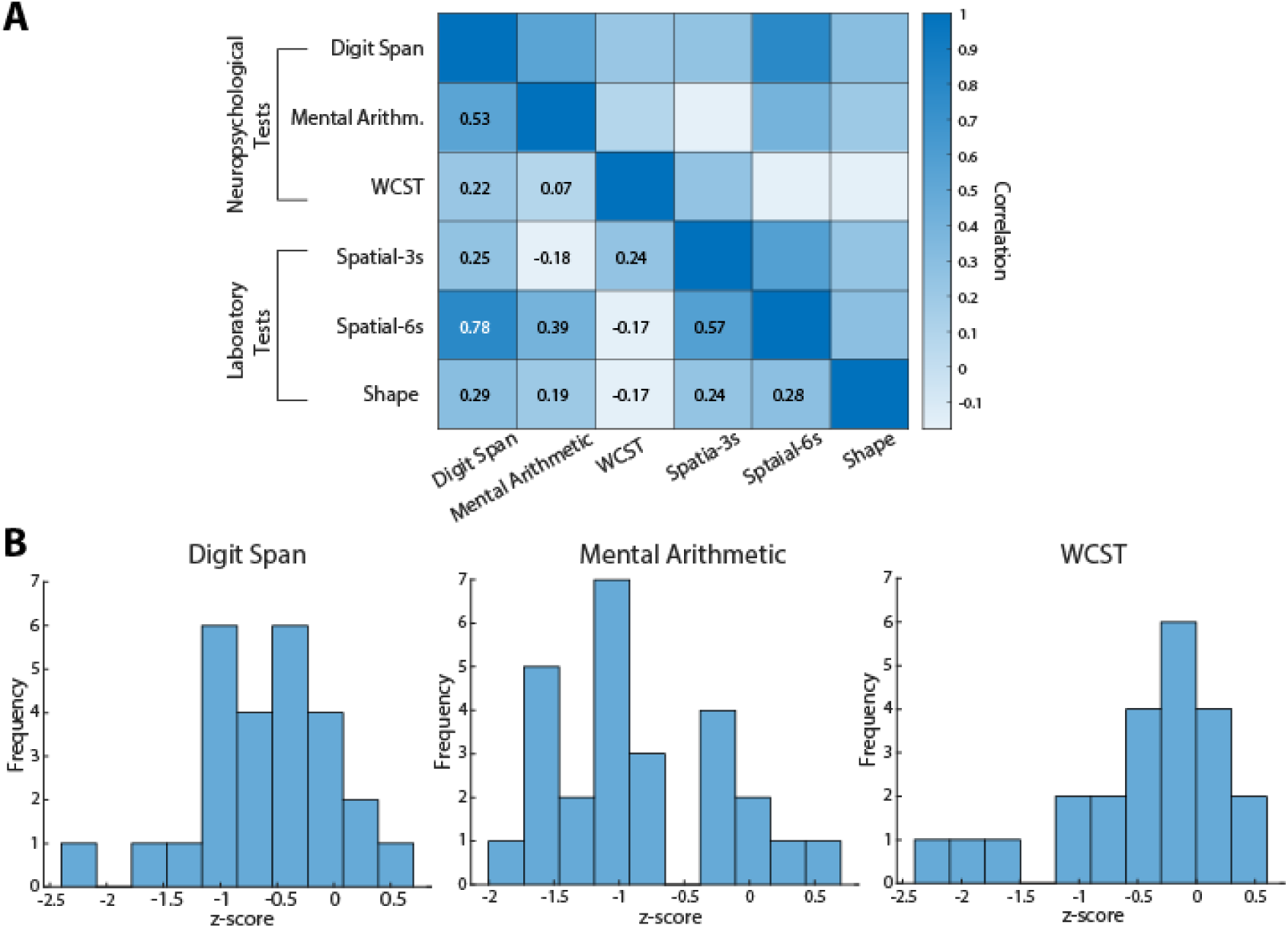
Performance in neuropsychological and laboratory tasks. A. Matrix representing correlation between scores in different neuropsychological (digit span, mental arithmetic, and Wisconsin Card Sorting Test – WCST) and laboratory tests (3s spatial working memory task; 6s spatial working memory task; shape match-nonmatch task). B. Score distribution from all subjects for the three neuropsychological tasks used: digit span task (n=26 subjects); mental arithmetic test (n=26 subjects) and Wisconsin card sorting task (n=23 subjects).

### Characteristics of LFP activity

LFP data were recorded from implanted intracranial electrodes which sampled a variety of cortical and subcortical regions. To simplify visualization and analysis we grouped recordings into 6 regions: mesial temporal, cingulate, lateral temporal, occipital, parietal and prefrontal regions (see Methods). Electrodes were excluded from data analysis if they were located in the patient’s seizure onset zone based on review of seizures by the clinical team.

Spectral power was normalized to mean power in each frequency bin during the whole trial. Time-resolved spectrograms were then calculated and averaged across contacts within each region. As we reported previously, for a partially overlapping dataset with that of the current study (Singh et al., 2024), we found neural responses from all six composite regions during both the spatial and shape tasks. Specifically, there was a brain-wide broadband signal during the cue presentation periods of the tasks (Fig. 3A). Additionally, there was a distinct trough in beta activity directly following cue onset. The magnitude of these responses varied between regions with an area-specific pattern (Fig. 3B and Supplementary Fig. S1).

**Figure 3.**
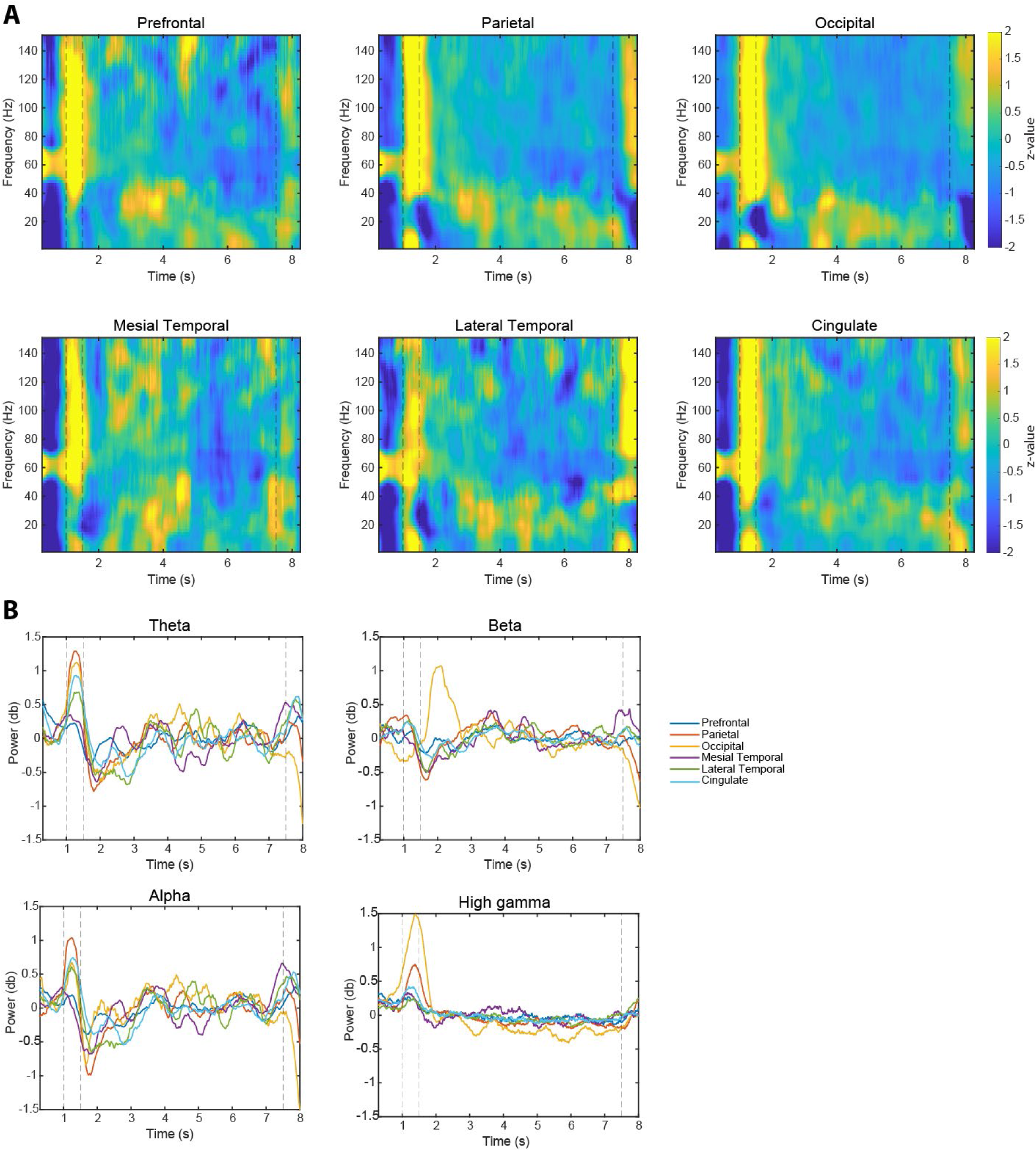
Time course of LFP spectral power. A. Mean, normalized LFP power in the 6s spatial working memory task. For each frequency (row) power has been normalized along the time axis to produce a z-value, illustrated by the color of each cell. B. Time course of power (relative to trial-average), plotted separately for the theta, beta, alpha, and high-gamma bands.

First, to identify electrode contacts that were task responsive during the working memory tasks, we used paired t-tests to identify contacts in which LFP activity in any of the studied frequency bands, specifically the theta (4-8 Hz), alpha (8-16 Hz), beta (16-40 Hz), and high-gamma range (70-150 Hz) activity differed significantly (p < 0.025) during one of the task epochs compared to the pre-cue fixation period. Of the 5406 total contacts between the three tasks, 2856 were found to be task-responsive in this manner. Next, we examined correlation between task-induced LFP activity in four bands of interest (theta, alpha, beta, and high gamma) in task responsive contacts and performance in the neuropsychological measures (as illustrated in Supplementary Fig. S2).

### Frequency and timing of LFP activity predictive of neuropsychological test performance

Considering the known activation of the prefrontal cortex, in animal studies and humans, during working memory (Constantinidis and Klingberg, 2016; Haller et al., 2018), we hypothesized that decreased prefrontal activation during the laboratory tests, using the high-frequency amplitude of the LFP as a proxy for neuronal firing, would be predictive of impaired working memory measured by neuropsychological tests. This prediction did not prove generally correct.

We tested the relationship between neural power and neuropsychological test scores on a sliding window through each task and performed cluster-based permutation testing to identify periods of significant relationships. Examining the correlation between high-gamma power and performance in neuropsychological tasks, the most notable finding was a negative correlation between LFP power during or after the stimulus presentations and performance in the neuropsychology tasks. This effect reached statistical significance for the correlation between high-gamma power following the second stimulus presentation in the shape working memory task and digit span performance (significant cluster: time= 4.5 – 5.3s, cluster p=0.019; Fig. 4A). A significant negative correlation was also observed between performance of the WCST task and LFP high-gamma power following the cue presentation in the 6s spatial task (significant cluster: time=1.6 – 2.2s, cluster p=0.002; Fig. 4C). A similar but non-significant negative correlation between high-gamma LFP power after the cue presentation in the 3s spatial task (peaking at the 1.9s time point in Fig. 4B) and performance of the digit span task was also observed. In other words, subjects who performed more poorly in the working memory neuropsychological tasks, tended to generate relatively more high-gamma power after the stimulus presentations.

**Figure 4.**
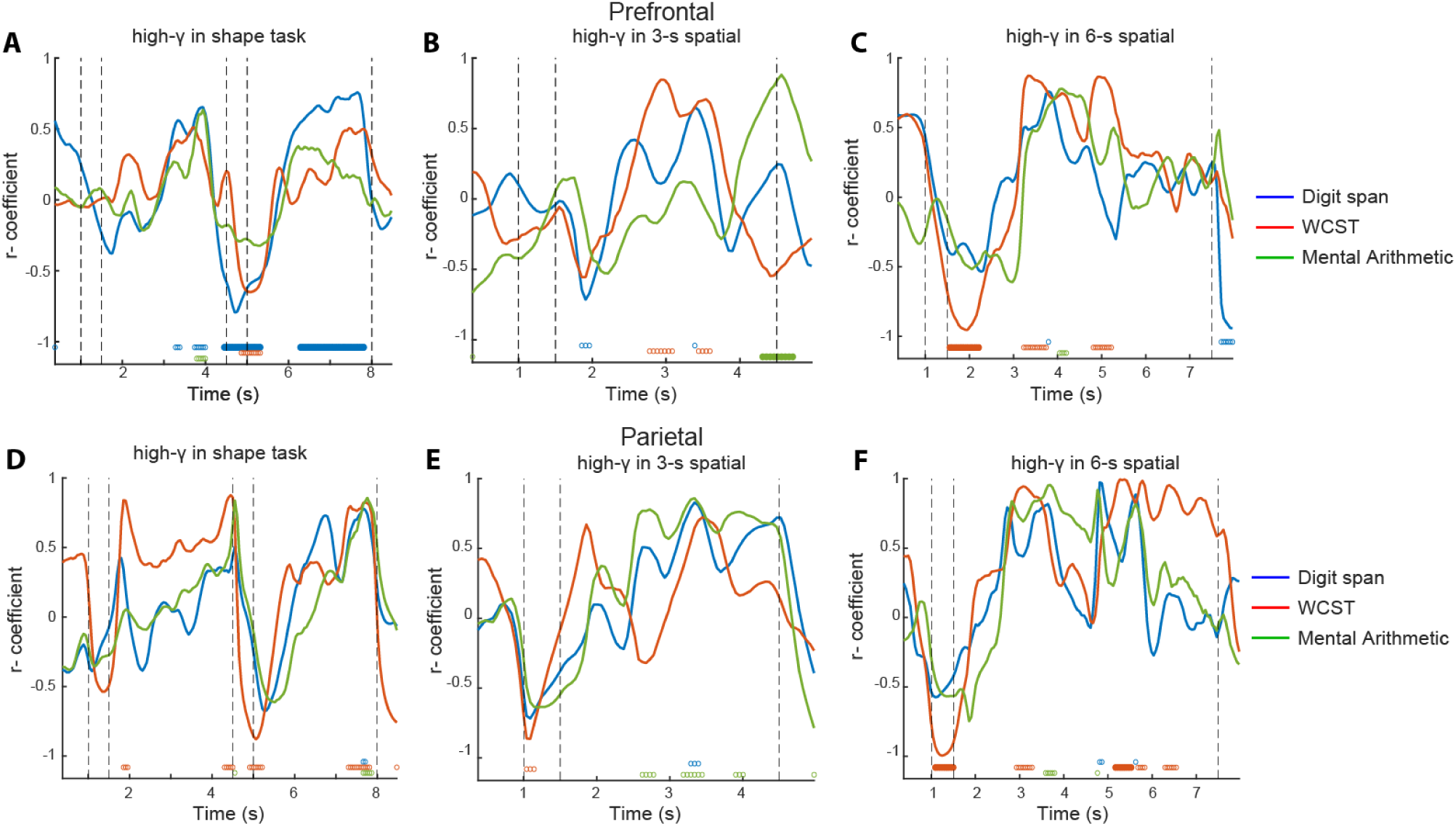
Correlation between high-gamma power and neuropsychological working memory test measure performance. In each plot, Pearson correlation coefficients between high-gamma activity observed in each time point of the three laboratory tasks and performance in a neuropsychological instrument are plotted. A. Correlation between prefrontal power during the shape working memory task and performance in the digit span, WCST, and mental arithmetic tasks (n=14 subjects, 184 contacts for digit span and mental arithmetic; n=13 subjects, 178 contacts for WCST). Open circles represent time points of significant relationships (p<0.05); larger filled circles mark significant clusters. B. Correlation for prefrontal high gamma in the 3s delay spatial task (n=10 subjects, 108 contacts for digit span and mental arithmetic; n=9 subjects, 102 contacts for WCST). C. Correlation for prefrontal high gamma in the 6s delay spatial task (n=7 subjects, 72 contacts for all tests). D. Correlation between high-gamma parietal power during the shape working memory task and performance of the digit span, WCST, and mental arithmetic tasks (n=7 subjects, 69 contacts). E. Correlation for parietal high gamma in the 3s delay spatial task (n=7 subjects, 63 contacts). F. Correlation for parietal high-gamma in the 6s delay spatial task (n=5 subjects, 48 contacts).

We sought to test whether this finding was consistent for the parietal cortex, another area that has been implicated in visual working memory and cognitive tasks (Hart and Huk, 2020; Qi et al., 2015; Xu, 2023). The results were similar to the prefrontal cortex (Fig. 4D-F). We found a negative correlation between high-gamma power following the second stimulus in the shape working memory task and WCST performance. Similarly, a negative correlation was present between performance of the WCST task and high-gamma LFP power during the cue presentation of the 6s spatial task (significant cluster: time=0.9 – 1.5s, p<0.001 in Fig. 4F).

On the other hand, in better agreement with our initial hypothesis, brief episodes of positive correlation between high-gamma power and neuropsychological test performance were observed during the delay periods of the laboratory tasks. Significant positive correlation was thus present between high gamma in the second half of the delay periods of the shape task and performance in the digit span task in (significant cluster: time= 6.4 – 7.8s, p=0.002 Fig. 4A); and high-gamma in the middle part of the delay period of the 6s tasks and WCST performance (significant cluster: time= 3.2 – 3.7s, p = 0.042 Fig. 4C). A similar but non-significant relationship was observed between high gamma power during the middle of the delay period of the 3s task and WCST performance (Fig. 4B). Similar positive correlation between high gamma in the delay period of the laboratory tasks and neuropsychological task performance was also observed for the parietal cortex. For example, a negative correlation was present between high gamma the following cue presentation during the 6s task and WCST performance (significant cluster: time=1.1 – 1.5s, p<0.001 Fig. 4F), and a positive correlation was present between high gamma in the middle of the delay of the 6s spatial tasks and WCST performance (significant cluster: time= 5.17 – 5.53, p<0.001, Fig. 4F). Similar but non-significant patterns were observed during the shape task and the 3s spatial task (Fig. 4D-E).

We performed a similar analysis of other frequency bands. In general, we found the opposite trend in theta- and alpha-frequency ranges compared to high-frequency amplitude (Supplementary Fig. S3A-F). Negative correlations were present between prefrontal theta- and alpha-power in the delay period with performance of the neuropsychological tasks (e.g. prefrontal theta during the delay period of the 3 second delayed response task vs WCST: time= 4.4 – 5.0s, p=0.005; prefrontal alpha during the delay period of the 3 second delayed response task vs WCST: t=4.4 – 5.0s, p=0.012). The relationship between power in the beta-range in the prefrontal and parietal cortex and neuropsychological performance was less consistent across tasks (Fig. S3-G-I). Results in other brain areas did not reveal as strong or systematic relationships between power and performance of the cognitive tasks (Fig. S4).

We performed additional time-frequency analysis to find significant relationships between induced power and executive function performance on the entire spectrogram (Fig. 5 and Supplementary Fig. S5). This analysis confirmed the findings from our primary analysis. High-frequency power around the times of stimulus presentations was negatively correlated with performance across neuropsychological tasks, and in both the prefrontal (Fig. 5A-C) and parietal cortex (Fig. 5D-F). Clusters of positive correlation between high-frequency power and neuropsychological test performance were encountered mostly during the delay period.

**Figure 5.**
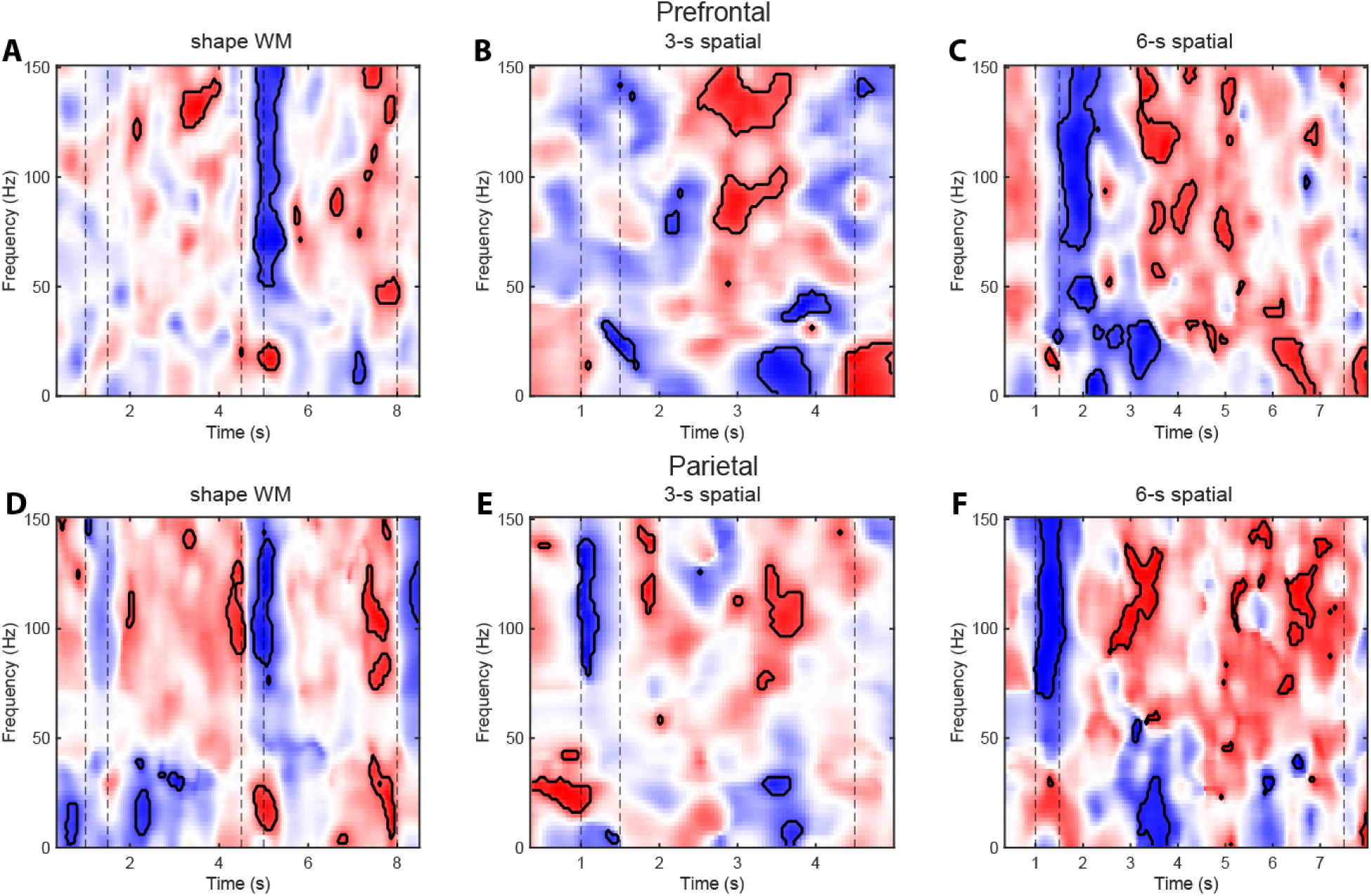
Time-frequency power correlation of prefrontal activity to WCST performance. Clusters are pixels with significant positive (red) or negative (blue) correlation (p < 0.05). A**-**C Correlation between prefrontal activity and performance in the shape working memory task (A); 3s spatial task (B); 6s spatial task (C). D-E Correlation between parietal activity and performance in the shape working memory task (D); 3s spatial task (E); 6s spatial task (F).

### Relationship between LFP activity and cognitive function

The comprehensive overview provided by the time-resolved, pairwise analyses above revealed similar temporal evolutions of correlation with LFP power by all three instrument scores. This indicates a potential stereotypical temporal pattern of which the activation magnitude could explain the cognitive function difference across subjects. To uncover such patterns, we conducted principal component analysis (PCA) and identified the first principal component (PC1) which captured most of the between-subject variance (Fig. 6). We focused specifically on prefrontal alpha and high gamma activity, since these bands both produced clusters that were significant when applying a correction for multiple comparisons, and our results suggested that these were not positively correlated with each other. Indeed, the uncovered PC1 mimicked the temporal pattern of the correlations. Both alpha and high gamma bands’ variance across subjects was largely captured by the depth of modulation between the cue and delay periods in all three tasks except for alpha band in the 3s spatial task (Fig. 6B). The percentage of variance explained by the first alpha PC was 32.9, 54.5, and 45.5, for the shape, 3s spatial, and 6s spatial tasks, respectively, while the first high-gamma PC explained 34.8, 34.1, and 42.0% of the variance, respectively.

**Figure 6.**
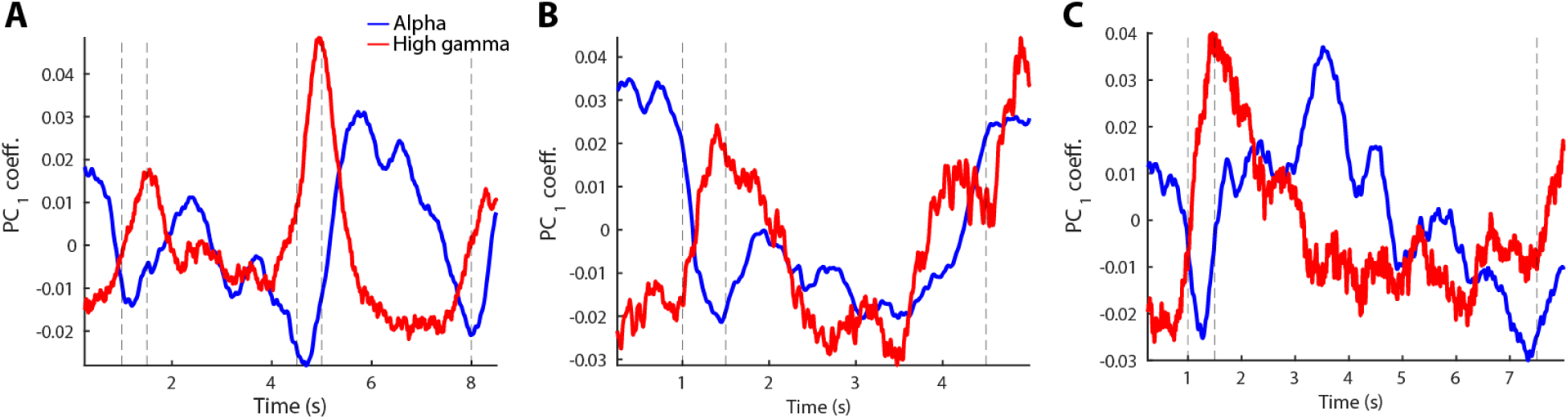
Principal Component Analysis. First principal component for alpha and high-gamma bands from the prefrontal cortex, of the shape task (A), 3s spatial task (B) and 6s spatial task (C).

Quantifying the amount of task activation of each subject as the PC1 score led us to develop a composite score combining our neuropsychological instruments that assessed several related, but not fully overlapping cognitive functions, that can be most robustly informed by the neural features recorded during our laboratory tasks. To this end, we used the subjects’ PC1 scores for each band as variables for canonical correlation analysis (CCA). We defined an LFP-power canonical variate U, incorporating scores for alpha and gamma power, and a cognitive canonical variate V, using digit span, WCST, and mental arithmetic scores. We then performed CCA to find the largest correlation between U and V and compared it against a null distribution produced by shuffling subject labels.

This analysis revealed that linear combination of the prefrontal alpha and gamma power was significantly correlated with the compound neuropsychological instrument score (r=0.87, permutation-test p=0.029, Fig. 7A, Supplementary Fig. S6A). The same analyses were not significant for the spatial tasks (r=0.897, permutation-test p=0.19 for 3s spatial task; r=0.981, permutation test p=0.11 for 6s spatial task, Fig. 7B-C, Supplementary Fig. S6B-C). To identify which predictors were the most influential in the relationship, we calculated the normalized magnitudes of each predictor’s loading on its respective canonical variate and determined the cosine similarity between our observed loadings and a unit vector and compared to a null distribution.

**Figure 7.**
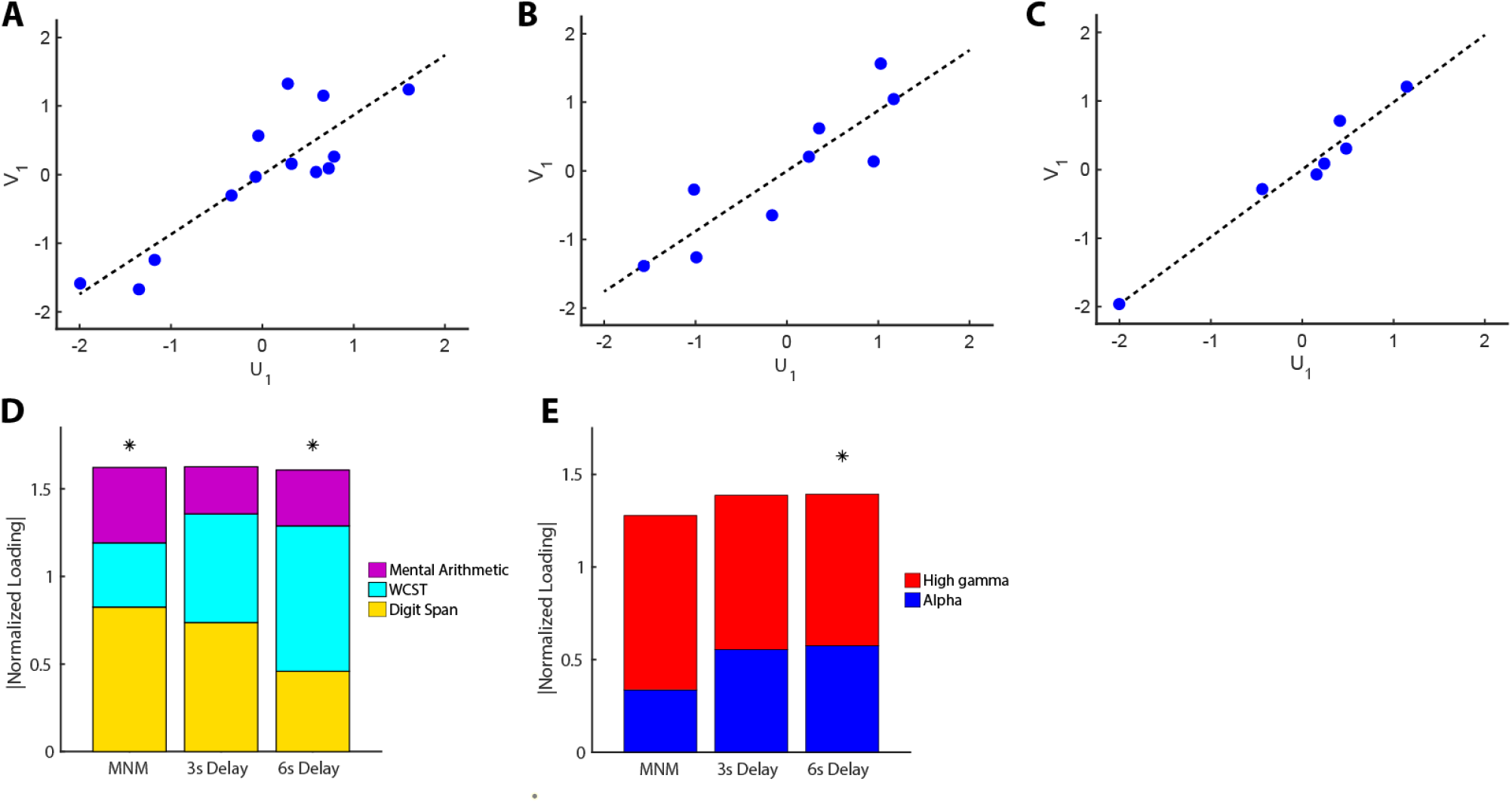
Canonical Correlation Analysis. A-C. Correlation between first cognitive canonical variate, U_1_, and first neural canonical variate, V_1_ for shape task (A), 3s spatial task (B) and 6s spatial task (C). **D** Magnitude of normalized loadings of alpha and gamma frequency bands on the neural canonical variate for each task. E) Magnitude of normalized loadings of task scores on the cognitive canonical variate. For D-E, asterisks represent loadings that significantly deviated from a unit vector.

Neuropsychological instrument score loadings were significantly different from equal loadings for the shape and 6s spatial task (p=0.041 for shape task, p=0.038 for the 6s spatial task), for which digit span and WCST were collectively more loaded than mental arithmetic (Fig. 7D). Loading difference did not reach significance for the 3s spatial task, although the overall pattern remained. Loadings for the neural canonical variate were significantly different from that of the unit vector for the 6s spatial task (p=6.7e-3) for which gamma power was more heavily loaded (Fig. 7A-B). This pattern remained for the other tasks, though their loadings were not significant different between bands. These results suggest that typical patterns of neural activity generated during performance of laboratory working memory tasks does capture cognitive capabilities measured independently by neuropsychological instruments, and combining distinct aspects of LFP activity can identify manifolds in the neuropsychological instrument space along which subject scores associate most closely with such measurable activities.

## DISCUSSION

Our results show that working memory ability, captured by neuropsychological instruments such as the digit span, mental arithmetic, and Wisconsin card sorting tasks, is associated with the activation of the prefrontal cortex during visual working memory laboratory tasks. We initially hypothesized that subjects with better working memory ability would generate higher activation of the prefrontal cortex. This hypothesis was spurred by findings in animal studies that have revealed higher levels of prefrontal activation particularly in the delay period of experimental conditions in which monkeys achieve higher performance (Kim and Shadlen, 1999; Qi et al., 2011); and in young adult animals compared to either juvenile or aged animals (Wang et al., 2011; Zhou et al., 2016). In the present study, there was some positive correlation between prefrontal high-gamma activity in the delay period of the task and neuropsychological test performance, however the effect was brief and did not span the entire delay period. Much more robust was a negative correlation between prefrontal and parietal high-gamma activity after the presentation of visual stimuli and working memory performance. This negative relationship was consistent for performance across different neuropsychological instruments, and across different laboratory tasks that were used to obtain the LFP signal. Activity in other frequency bands and in other brain areas was generally less consistent in predicting performance of neuropsychological tasks, though alpha generally moved in the opposite direction and a linear combination prefrontal alpha and high gamma power captured well a compound neuropsychological instrument score incorporating all instruments.

### Prefrontal activation as a marker of cognitive performance

Examples linking increased prefrontal activation with lower cognitive function have been provided by human imaging studies. An abnormally high prefrontal activation has been implicated in fMRI studies of aging and other pathological conditions, interpreted as a mechanism to compensate deficiency, or reflecting greater effort to complete a cognitive task (Grady, 2012). Some human imaging studies in healthy, young adults have also suggested decreases in Blood Oxygenation Level-Dependent (BOLD) activity after training that improves performance in working memory tasks, a phenomenon that has been interpreted as an increase in efficiency (Kuhn et al., 2013; Schneiders et al., 2011; Schneiders et al., 2012; Schweizer et al., 2013; Takeuchi et al., 2013). Our present results implicate increased prefrontal and parietal high-gamma power after the presentation of visual stimuli as the most reliable indicator of impaired working memory ability.

An important caveat for the interpretation of these results is that our analysis was based on LFP amplitude at each frequency normalized across the entire trial for each patient and electrode contact (Haller *et al*., 2018; Weber et al., 2023). It is possible that the absolute firing rate of neurons in the prefrontal cortex of high-performing patients achieved a higher firing rate during the task that was somewhat masked by a higher baseline firing rate as well. The relative power of the high-gamma component of the LFP relative to this baseline is still an important characteristic of neural activity that could serve as biomarker of working memory ability and executive function.

The presence of epilepsy in the subjects of our study is also likely associated with atypical brain activity and connectivity. Our sample’s mean neuropsychological performance was markedly below the population average. Finally, our comparisons involved fairly coarse brain areas, as we did not have sufficient power to investigate subdivisions of the prefrontal cortex that have been implicated in specific aspects of working memory and cognitive control (Badre and Nee, 2018; Miller et al., 2022). These limitations notwithstanding, our results demonstrate that it is possible to identify features of neural activity that are associated with working memory performance.

### Neural basis of working memory

The neural basis of working memory has been debated in recent years (Constantinidis *et al*., 2018; Lundqvist et al., 2018). Persistent activity selective for the spatial location of the remembered stimulus has been reported in neurophysiological studies of the prefrontal cortex in non-human primates performing working memory tasks (Funahashi et al., 1989). Error trials in the visually guided saccade task (similar to our spatial task in this study but involving an eye movement) are characterized by lower levels of delay period activity (Funahashi *et al*., 1989; Zhou et al., 2013). A near linear relationship between behavioral performance and persistent activity can be also revealed in tasks that modulate parametrically the discriminability of the remembered targets (Constantinidis et al., 2001) or across sessions grouped based on performance (Zhou *et al*., 2016). Computational models provide a detailed relationship between working memory performance and persistent activity; drifts in neuronal activity predict the endpoint of the saccade, the spatial location being recalled by the monkey (Barbosa et al., 2020; Wimmer et al., 2014). In these bump-attractor models, appearance of a stimulus generates activity which is maintained during the delay period but may drift with time.

Although this model emphasizes a direct relationship between firing rate and performance of working memory tasks, it is well understood that decrease in baseline firing rate may improve working memory performance (Tang et al., 2019). Other aspects of neuronal firing, such as correlated variability are also modulated with training that improves performance (Qi and Constantinidis, 2012) and could contribute to differences in performance but may not be directly visible at the level of LFP responses.

Alternative models have also been proposed that do not depend on persistent activity to represent information during the delay period of working memory tasks. The rhythmic-burst model depends on gamma band rhythmicity of prefrontal activity, instead. Gamma band rhythmicity in the prefrontal cortex is modulated during working memory, and scales with working memory load and other task parameters (Lundqvist et al., 2016; Bastos et al., 2018; Lundqvist et al., 2018b; Stanley et al., 2018; Wutz et al., 2018). A third class of models suggest that information can be decoded from the prefrontal cortex without generation of elevated persistent activity across the population, in an “activity-silent” manner (Stokes, 2015). Experimental studies have shown that the identity of a stimulus maintained in memory can be decoded from a population of neurons whose activity is not elevated above baseline levels of firing rate (Stokes et al., 2013). These results suggest that more refined analysis will be necessary to understand the neural processes that govern working memory ability in the human brain. Studies relying on single neuron activity in the human prefrontal cortex may ultimately resolve these questions. Our current results do demonstrate that levels of neuronal activation are not only relevant to the underlying neural mechanisms of working memory but could potentially inform future diagnostic and treatment methods for cognitive dysfunction.

## STAR METHODS

### RESOURCE AVAILABILITY

#### Lead Contacts

Further information and requests for resources and reagents should be directed to and will be fulfilled by the Lead Contacts, Dr. Sarah Bick and Christos Constantinidis.

#### Materials Availability

This study did not generate new unique reagents.

#### Data and Code Availability

- Data used for the analysis and figures will be deposited at Mendeley.com and be made publicly available as of the date of publication.
- This paper does not report original code.
- Any additional information required to reanalyze the data reported in this paper is available from the lead contact upon request.

### EXPERIMENTAL MODEL AND SUBJECT DETAILS

A total of 28 patients with medically intractable epilepsy who had stereo electro-encephalography (sEEG) electrodes implanted to localize their seizure focus were recruited in this study (10 male and 18 female). For the analysis of correlation between neuropsychological test measures and neural activity, we excluded 4 patients for whom neuropsychological assessments were performed too long (>24 months) before the neurophysiological recordings, one patient for which the LFP record was contaminated by high frequency noise and one patient for which neuropsychological test scores were not available. This resulted in a sample of 22 patients (9 male, 13 female) used for analysis. Following surgery, patients were admitted to the epilepsy monitoring unit and antiepileptic medications gradually weaned while patients were continuously monitored for seizures. All subjects gave informed consent for this study, which was approved by the Institutional Review Board of the Vanderbilt University Medical Center (Nashville, TN, IRB #211037).

## METHODS DETAILS

### Neuropsychological Instruments

Prior to sEEG surgery, patients completed a neuropsychological battery that included tests to assess their executive function ability, including the Wisconsin Card Sorting Test 64 Card Version (WCST-64) and the Weschler Adult Intelligence Scale Fourth edition (WAIS-IV). We focused on tests most relevant to working memory. Other tasks, not analyzed here, included measures evaluating language, visual perception, memory, and sensorimotor domains. From the WCST-64, we focused on preservative errors and number of categories completed. From the WAIS-IV, we specifically looked at the mental arithmetic and digit span task performance (total score of forward, backward, and sequencing digit span). Raw test scores were converted to norm-referenced z-scores based on the general population using psychometric conversion tables. Scores from each task were averaged for a subject to obtain an overall WCST and Digit Span performance calculations.

### Laboratory Tasks

Participants performed visual working memory tasks using a portable tablet with a 26.0 cm-by-17.3 cm display and stylus (Microsoft Surface, Redmond WA) in their hospital rooms, while LFPs were continuously recorded from sEEG electrodes. Task control was achieved in MATLAB 2021 (MathWorks, Natick, MA) using custom Psychophysics Toolbox, Version 3 (Brainard, 1997). At the beginning of each task epoch, the serial port triggers an Arduino Leonardo to send a TTL pulse as a timestamp to the trigger channel of the EEG amplifier (Appelhoff and Stenner, 2021).

Variants of a manual delayed response task (spatial task) and a shape match-nonmatch task (shape task) were used. In the spatial task (Fig.1A), a circle 2 cm in diameter appeared at the start of each trial in the center of the tablet screen and the subject moved the stylus into the circle to initiate the trial. After a 1 s “fixation” period, a second white circle of the same size appeared (cue) at a peripheral location 6.1 cm away from the center of the display for 0.5 s, after which only the center circle remained. This is followed by a 3 s or 6 s (delay) interval where, again, only the fixation point was displayed. After the delay period, the center circle disappeared, and the subject needed to drag the stylus across the screen into an invisible circle 2.4 cm in diameter centered at the remembered location of the cue. The stylus needed to enter the invisible circle within 2 s and stay inside for 0.5 s for the trial to be considered correct. A TTL pulse marked the beginning of each task event. Delay durations were blocked; trials with 3 s delays were performed first, and then the 6 s delay trials were presented. Cue locations in a block were evenly distributed across 360 degrees starting at 0 degrees. A typical block had 36 cue locations, each of which appeared once. Cue locations in completed trials were not repeated within a block.

In the shape task (Figure 1B), at the start of each trial, a circle appeared in the center of the tablet screen, and the subject moved the stylus into the circle to initiate the trial. The circle remained visible for a 1 s fixation period. Then a stimulus was presented for 0.5 s, comprising a white, convex polygon (cue), replacing the center circle. This is followed by a 3 s (delay) interval where, again, only the white circle was displayed. A second stimulus (sample) was subsequently shown for another 0.5 s, followed by a second 3 s delay period. At the end of the second delay period, the center circle disappeared, and two colored circles appeared at the top and bottom of the screen. The subject then needed to drag the stylus across the screen into either the green circle or red circle to indicate whether the two polygons were the same (match) or not (nonmatch), respectively. The polygons used had 3 to 6 vertices, did not extend beyond the extend of the center circle and no polygon was a rotated version of another. The red and green circle placement was randomly switched between the top and bottom locations on each trial. A total of eight shapes were used, followed by either a top and bottom location in match or nonmatch to create 32 trials. The circle sizes and the response window duration in the shape task were the same as those in the spatial task.

Before participants performed the full block of the visual working memory tasks, a member of the experimenter team explained and demonstrated the procedure. The participants then practiced a few trials to ensure they understand the experimental protocol before the block began. These practice trials were excluded from analysis.

### Electrode implantation and localization

Electrodes were implanted under general anesthesia with implant locations determined on an individual subject basis by clinical considerations related to hypotheses for site of seizure onset. Participants were implanted with multiple electrode shafts of 0.8-1.3 mm in diameter and containing 8-16 contacts, spaced between 2.5-4.3 mm center-to-center (PMT corporation, Chanhassen, MN; and AdTech Medical Instrument corporation, Oak Creek, WI). We identified the location of all contacts in each individual’s brain by superimposing the pre-surgical MRI and postsurgical CT images using the CRAnial Vault Explorer (CRAVE) software (D’Haese et al., 2012) and the FreeSurfer software package (http://surfer.nmr.mgh.harvard.edu). These software tools allowed us to determine coordinates of each contact, and to identify the anatomical location of each contact according to the Desikan-Killiani Atlas (Desikan et al., 2006) using FreeSurfer’s cortical parcellation and subcortical segmentation procedure (Fischl et al., 2002; Fischl et al., 2004). White matter contacts were excluded from analysis. Electrode contact localizations obtained in this manner were additionally verified by an epileptologist (SWR) based on visual inspection of the co-registered MRI, postsurgical CT and baseline EEG timeseries.

Electrode contacts were classified into six broad areas (Singh *et al*., 2024) defined as follows: a) Mesial Temporal region, which included contacts from the Amygdala, Hippocampus, and Mesial Temporal Cortex; b) Cingulate region, which included the anterior, mid and posterior cingulate; c) Lateral Temporal region, which included contacts in the Inferior Temporal Cortex and Superior and Middle Temporal gyrus and Insula; d) Occipital region, which included any contacts in the occipital lobe; e) Parietal region, which included contacts in the parietal lobe (typically in the posterior parietal cortex); f) Prefrontal region, which included contacts in any subdivision of the prefrontal cortex.

### LFP Recording, preprocessing and signal analysis

Local field potentials were recorded from the implanted sEEG electrodes and sampled at 512 Hz, using the Natus clinical data acquisition system (Natus, Middleton, WI). Reference electrodes were placed on the scalp. Task events were synchronized with the LFP recording with a TTL pulse that was generated by the tablet and was recorded in the data acquisition system. Task epochs (e.g., fixation, stimulus presentation, delay period) were aligned to different TTL pulses. LFP recordings were preprocessed by using custom MATLAB code in MATLAB R2022b (MathWorks) and the FieldTrip toolbox (Oostenveld et al., 2011). A bandpass filter between 0.5-200 Hz with a zero-phase sixth-order Butterworth filter were used on single-trial LFP traces. To remove 60 Hz power line noise and its harmonics, an infinite impulse response (IIR) Butterworth filter was applied. Electrodes were excluded from data analysis if they were not in gray matter or were determined to be within the patient’s seizure onset zone based on review of seizures by the clinical team and confirmed by a clinical epileptologist (SWR).

To minimize the influence of artifact waveforms, such as those caused by interictal epileptiform discharges and movement, we used an iterative procedure to objectively remove outlier trials from each electrode’s sample pool. Within a contact channel, trials were removed if their LFP trace had a kurtosis >2.8 or a variance >2.2 SD from the persistent sample mean (Sheehan et al., 2018). Further, to identify trials with frequency-specific artifacts, we calculated the Euclidean distance of each trial’s induced spectrogram to the mean spectrogram across trials. Trials with a Euclidean distance >2.2 SD from the persistent sample mean were removed. Finally, we identified task-responsive electrodes by testing for spectral power changes in any of four frequency bands (theta 4-8 Hz, alpha 8-16 Hz, beta 16-40 Hz, and high gamma 70-150 Hz) during any of the laboratory tasks. A paired t-test was used to determine if spectral power was significantly different during either the cue or delay period compared to the baseline fixation period. For the shape task, both cue periods and both delay periods were considered. Contacts showing significant power changes (p < 0.025) in any of the frequency bands during any task epoch of any laboratory task were considered task responsive. This test was blind with respect to the direction of change from baseline. Only responsive contacts were used for further analysis.

The Chronux package (Bokil et al., 2010) was used for time-frequency analysis. We used a multi-taper method to perform a power spectrum analysis of LFPs. The spectrogram of each single trial between 0.5 and 150 Hz was computed with 8 tapers in 500 ms time windows; the spectrograms were estimated with a temporal resolution of 2 ms. We also used the mean filter corresponding to 2 Hz and 2 ms for smoothing the spectrogram of each single trial. In all our analyses, we relied on induced power of the LFP, which is computed by first performing a power computation in each trial and then averaging power across trials. Power in each frequency band was expressed relative to the mean power recorded during the entire trial beginning at -500 ms before the first cue onset and ending 500 ms after the response. We constructed time-resolved plots (spectrograms) by dividing the power of the signal by the mean power at each frequency. We calculated each contact’s overall power in frequency bands by averaging the contact’s mean spectrogram over frequency ranges for each band of interest.

### Statistical Analysis

We first calculated the time-resolved correlation between subjects’ neuropsychological test scores and LFP power from the previously defined frequency bands. In each region of interest, we calculated the Pearson’s correlation coefficient between the log-scaled mean power and test score on a 250 ms sliding window with a 50 ms step through the duration of each task. Subjects with fewer than three responsive contacts in any given area were excluded from correlation analysis for that area. We then calculated the Pearson’s correlation coefficient between region power in each frequency band and Neuropsychological test scores. We used a cluster-based permutation test to identify time periods of significant relationships while controlling for family-wise error rate (Oostenveld et al 2011). In each region of interest, we first performed a simple linear regression between test score and log-scaled mean power at each time point using a 250 ms window with a 50 ms step. Time bins yielding coefficients with a significant t-statistic based on a two-tailed t-distribution at α = 0.05 formed significant clusters. The t-statistics from each time bin were summed in each significant cluster to obtain a cluster-level statistic. Scores were then randomly shuffled, and the process was repeated 1000 times. The largest cluster-level statistic from each permutation was used to form a null distribution. Observed clusters were identified as significant if their cluster-level statistic fell in the 95^th^ percentile of the null distribution. To make the direction and magnitude of relationships more interpretable, regression coefficients were transformed to Pearson’s correlation coefficients for visualization.

To get a broader idea of the relationship between frequency power and test score over time, we performed time-frequency-resolved correlation analysis using similar cluster-based permutation testing. We first performed simple linear regression between test score and log-scaled mean power at each time point using a 250 ms window and a 50 ms step along frequency bands 2 Hz in width.

We conducted canonical correlation analysis to determine whether neural features during our working memory tasks could be robustly related to cognitive function as measured by performance on our Neuropsychological instruments. We performed this analysis in subjects with scores available for all three neuropsychological instruments (n=13, n=9, and n=7 tested with the shape, 3s delay, and 6s delay spatial tasks, respectively). First, for each task, we performed principal component analysis (PCA) on the alpha and high-gamma traces from all subjects and extracted the first PC for each band. We then calculated subjects’ scores corresponding to the first PC, which represent their contribution to the observed typical pattern. Subjects scores for the alpha and beta bands were used as the neural canonical variate, V. For the cognitive score canonical variate, V, we used digit span, WCST, and mental arithmetic scores. We then performed simple linear regression between the first U and first V to obtain the correlation coefficient r_obs_. Because CCA with small sample sizes can produce inflated r values (Helmer et al., 2024), we compared r_obs_ to a null distribution that was made by randomly shuffling the order of the scores, while keeping the scores aligned relative to one another, 1000 times to determine the probability of obtaining a correlation coefficient of r_obs_ or greater by chance. We deemed r_obs_ significant if it’s p-value relative to the null distribution was <0.05.

To determine the relative contribution of each predictor to the observed relationships, we calculated their loadings on the respective canonical variate. For instance, we calculated the correlation coefficients between digit span, WCST, and mental arithmetic scores and the cognitive canonical variate to obtain each score’s loading. Loadings were then normalized to unit length and compared to a unit vector representing equal weight from each predictor using absolute value of cosine similarity. Absolute cosine similarity between two vectors can range from 0 to 1 where 1 represents equal vectors and 0 represents orthogonal vectors. The absolute cosine similarity between normalized loadings and the unit vector were calculated after shuffling scores 1000 times. Observed loadings were identified as significant if they were in the bottom 5^th^ percentile of similarity; that is, if the probability of finding a less evenly distributed vector of loadings was less than 0.05.

## ACKNOWLEDGEMENTS

Supported by NIH award number R01 EY017077 (CC), K23 AG072030 (SWR), and K12 NS080223 (SKB). We wish to thank Chrissy Suell and Kayla Yetman for technical help.

## AUTHOR CONTRIBUTIONS

CC, DJE, SWR and SKB designed the experiment. AJ, BS, ZW, MLJ, and CC collected data. AJ, BS, ZW, HQ performed data analyses. AJ, KD, ZQ, CC, SWR, SKB interpreted data. CC and AJ wrote the paper with input from all authors.

## SUPPLEMENTARY TABLES

**Supplementary Table 1.**
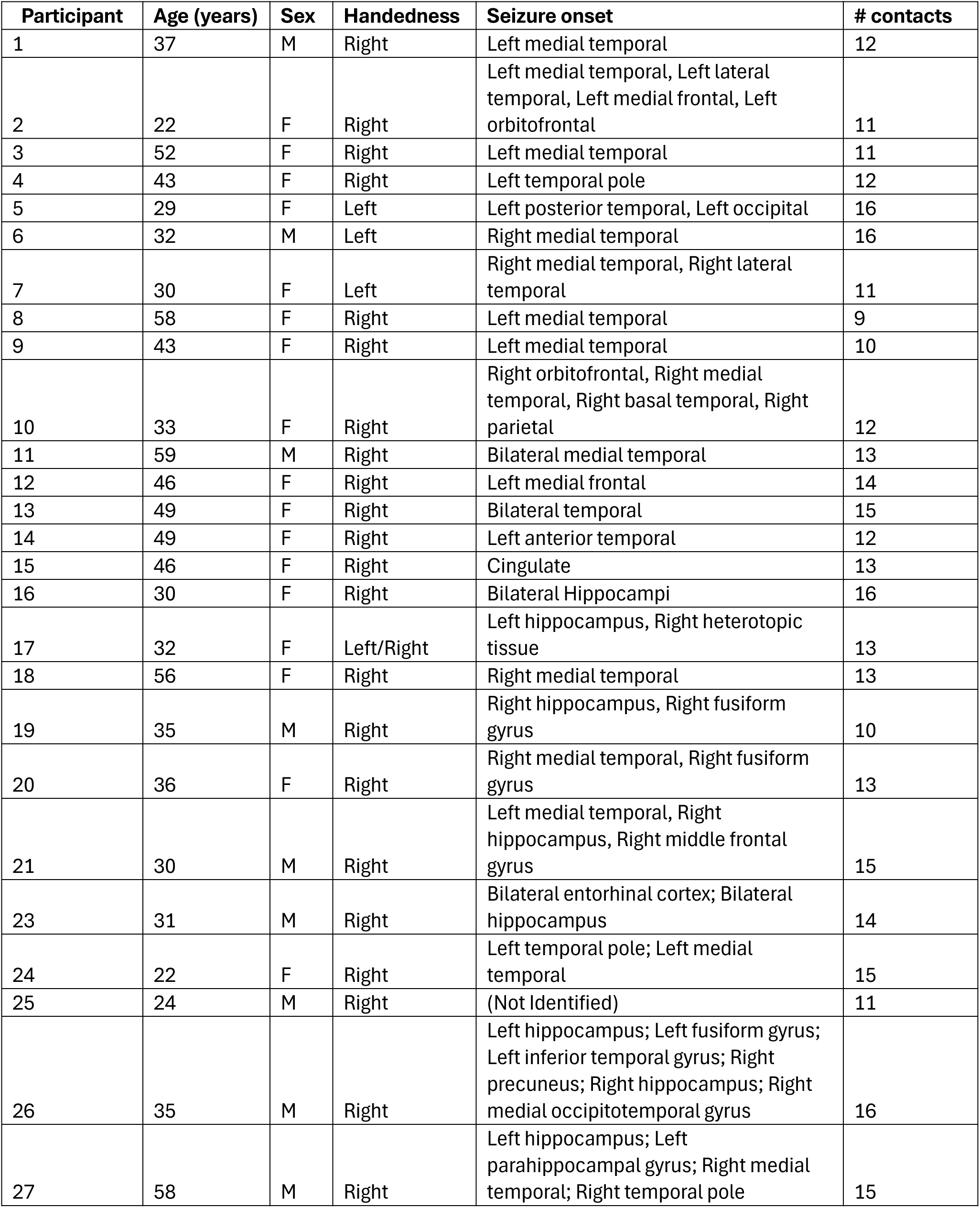
Patient demographic characteristics. Table reports age, sex, handedness, and seizure onset, as this was determined clinically, for study participants as well as number of electrode contacts used for analysis.

| Participant | Age (years) | Sex | Handedness | Seizure onset | # contacts |
| --- | --- | --- | --- | --- | --- |
| 1 | 37 | M | Right | Left medial temporal | 12 |
| 2 | 22 | F | Right | Left medial temporal, Left lateral temporal, Left medial frontal, Left orbitofrontal | 11 |
| 3 | 52 | F | Right | Left medial temporal | 11 |
| 4 | 43 | F | Right | Left temporal pole | 12 |
| 5 | 29 | F | Left | Left posterior temporal, Left occipital | 16 |
| 6 | 32 | M | Left | Right medial temporal | 16 |
| 7 | 30 | F | Left | Right medial temporal, Right lateral temporal | 11 |
| 8 | 58 | F | Right | Left medial temporal | 9 |
| 9 | 43 | F | Right | Left medial temporal | 10 |
| 10 | 33 | F | Right | Right orbitofrontal, Right medial temporal, Right basal temporal, Right parietal | 12 |
| 11 | 59 | M | Right | Bilateral medial temporal | 13 |
| 12 | 46 | F | Right | Left medial frontal | 14 |
| 13 | 49 | F | Right | Bilateral temporal | 15 |
| 14 | 49 | F | Right | Left anterior temporal | 12 |
| 15 | 46 | F | Right | Cingulate | 13 |
| 16 | 30 | F | Right | Bilateral Hippocampi | 16 |
| 17 | 32 | F | Left/Right | Left hippocampus, Right heterotopic tissue | 13 |
| 18 | 56 | F | Right | Right medial temporal | 13 |
| 19 | 35 | M | Right | Right hippocampus, Right fusiform gyrus | 10 |
| 20 | 36 | F | Right | Right medial temporal, Right fusiform gyrus | 13 |
| 21 | 30 | M | Right | Left medial temporal, Right hippocampus, Right middle frontal gyrus | 15 |
| 23 | 31 | M | Right | Bilateral entorhinal cortex; Bilateral hippocampus | 14 |
| 24 | 22 | F | Right | Left temporal pole; Left medial temporal | 15 |
| 25 | 24 | M | Right | (Not Identified) | 11 |
| 26 | 35 | M | Right | Left hippocampus; Left fusiform gyrus; Left inferior temporal gyrus; Right precuneus; Right hippocampus; Right medial occipitotemporal gyrus | 16 |
| 27 | 58 | M | Right | Left hippocampus; Left parahippocampal gyrus; Right medial temporal; Right temporal pole | 15 |

## SUPPLEMENTARY FIGURES

**Supplementary Figure S1.**
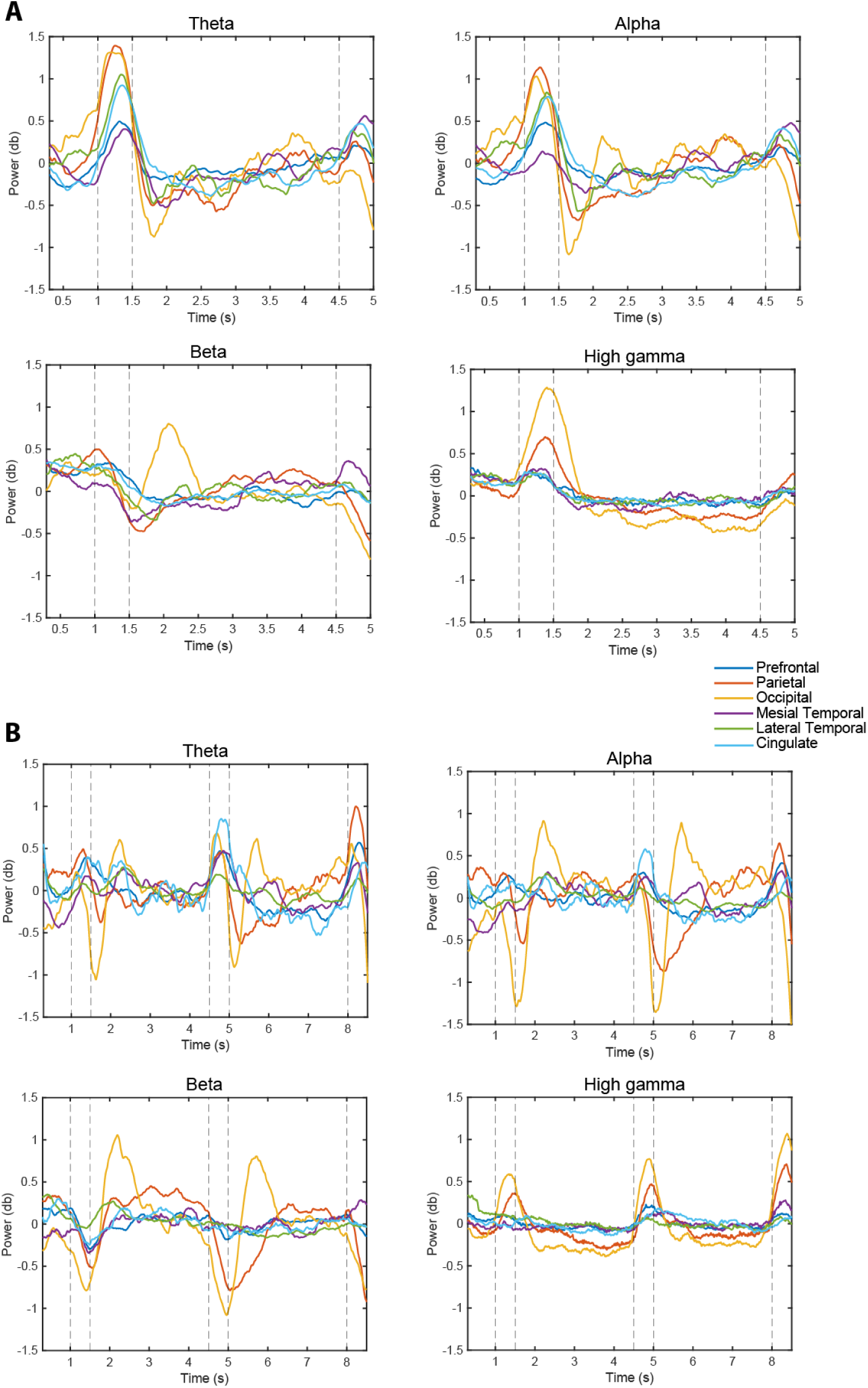
Time course of LFP spectral power in the 6s spatial and shape working memory tasks. A. Power relative to a trial-average baseline as function of time in the 3s spatial task, plotted separately for the theta, beta, alpha, and high-gamma bands, for each of the six brain regions. B. Power in the shape match-nonmatch task. Conventions are the same as in Fig. 3B.

**Supplementary Figure S2.**
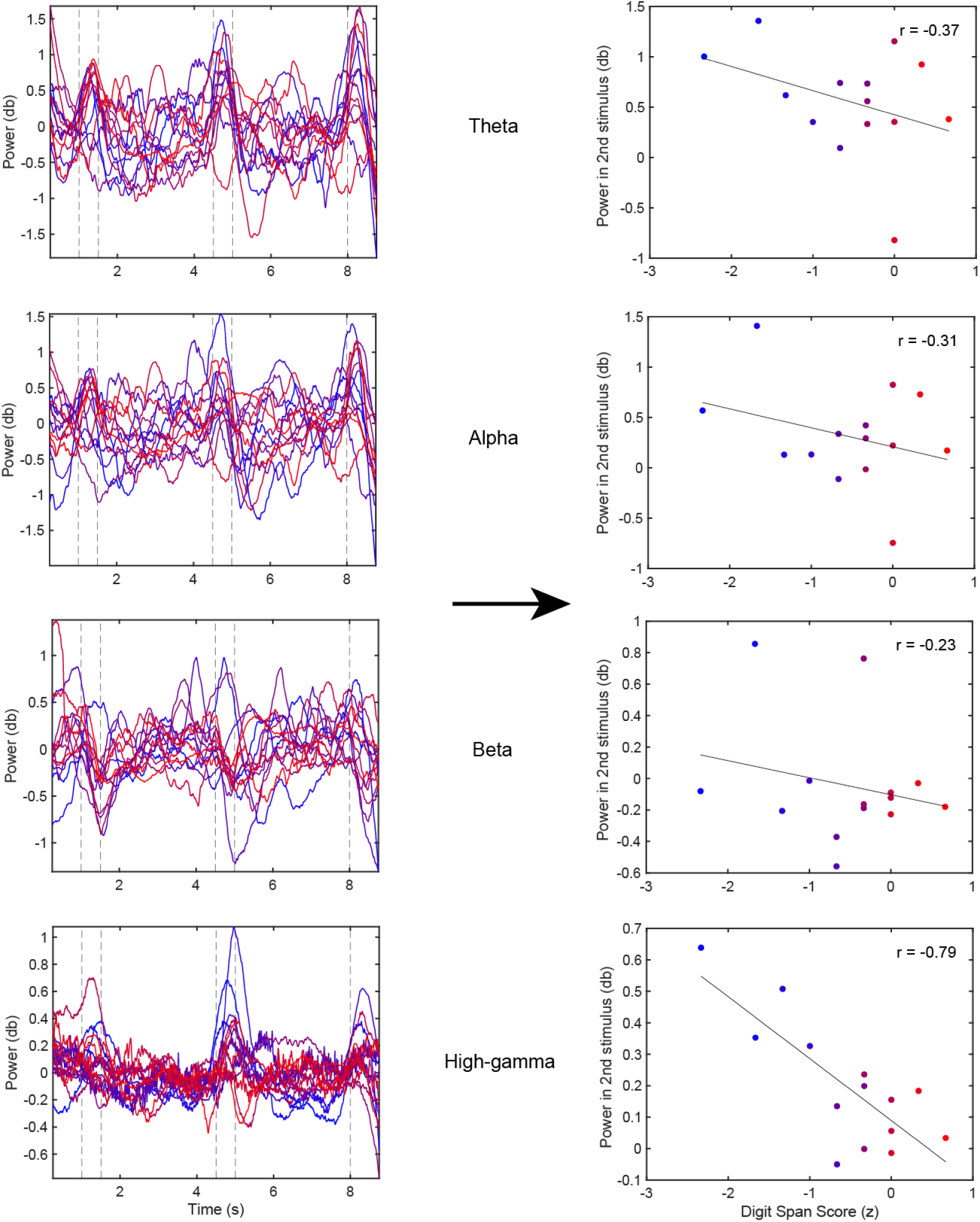
LFP power – performance correlation. Left, Power recorded during the shape task is plotted as a function of time, for each subject. Color of each trace represents the performance of each subject in the digit span task (red, higher performance; blue, lower performance). Right, Power during the second stimulus period is plotted as a function of performance in the digit span task. Each point represents one subject, color coded in the same fashion as in the left plots.

**Supplementary Figure S3.**
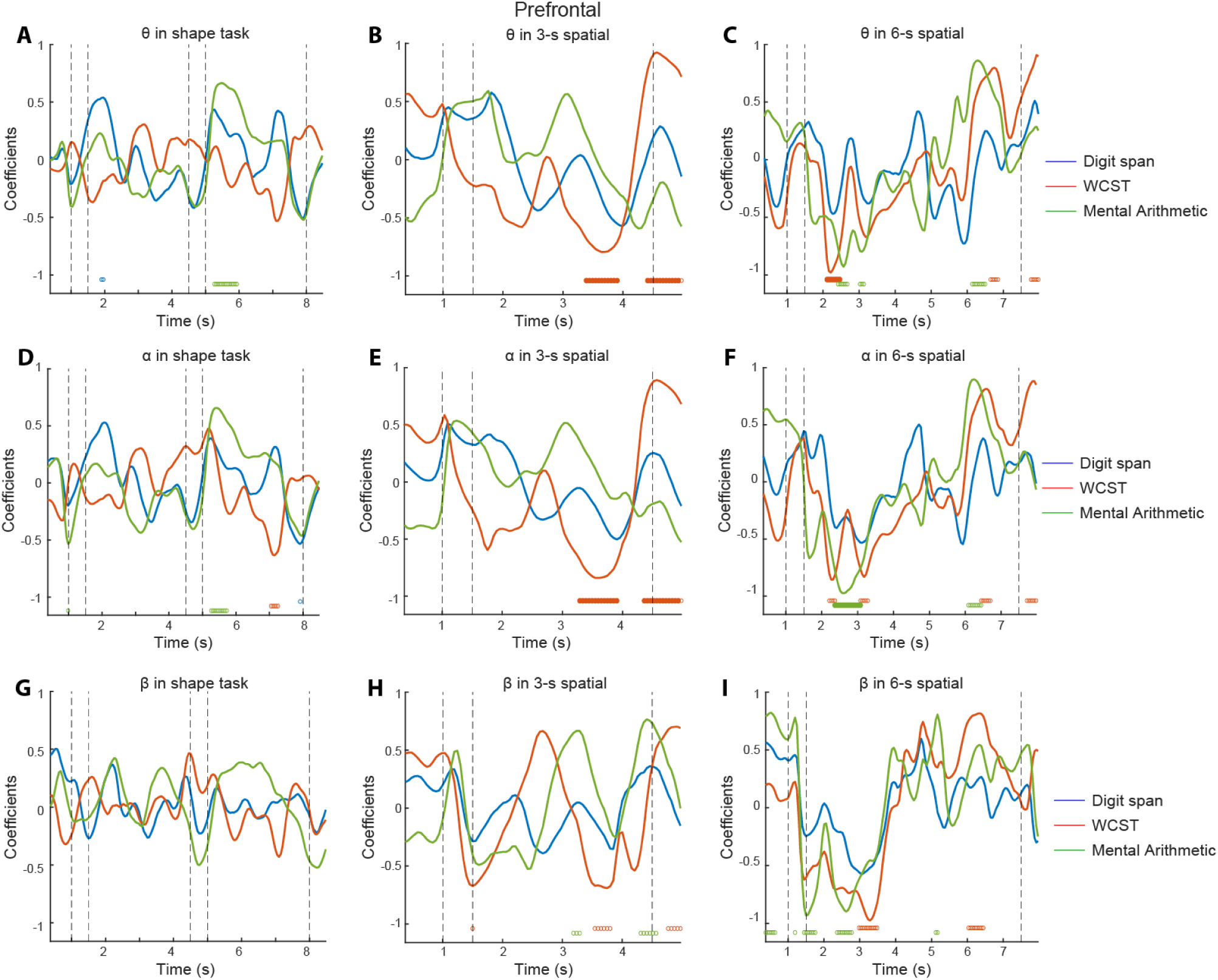
Correlation between lower-frequency bands of neural activity and neuropsychological test performance. A-C. Correlation between prefrontal theta power and performance of each of the neuropsychological tasks. A. Shape working memory task (n=14 subjects, 184 contacts). B. 3s delay spatial task (n=10 subjects, 108 contacts). C. 6s delay spatial task (7 subjects, 72 contacts). D-F. Correlation between prefrontal alpha power and performance. G-I. Correlation between beta power and performance. Conventions are the same as in Fig. 4.

**Supplementary Figure S4.**
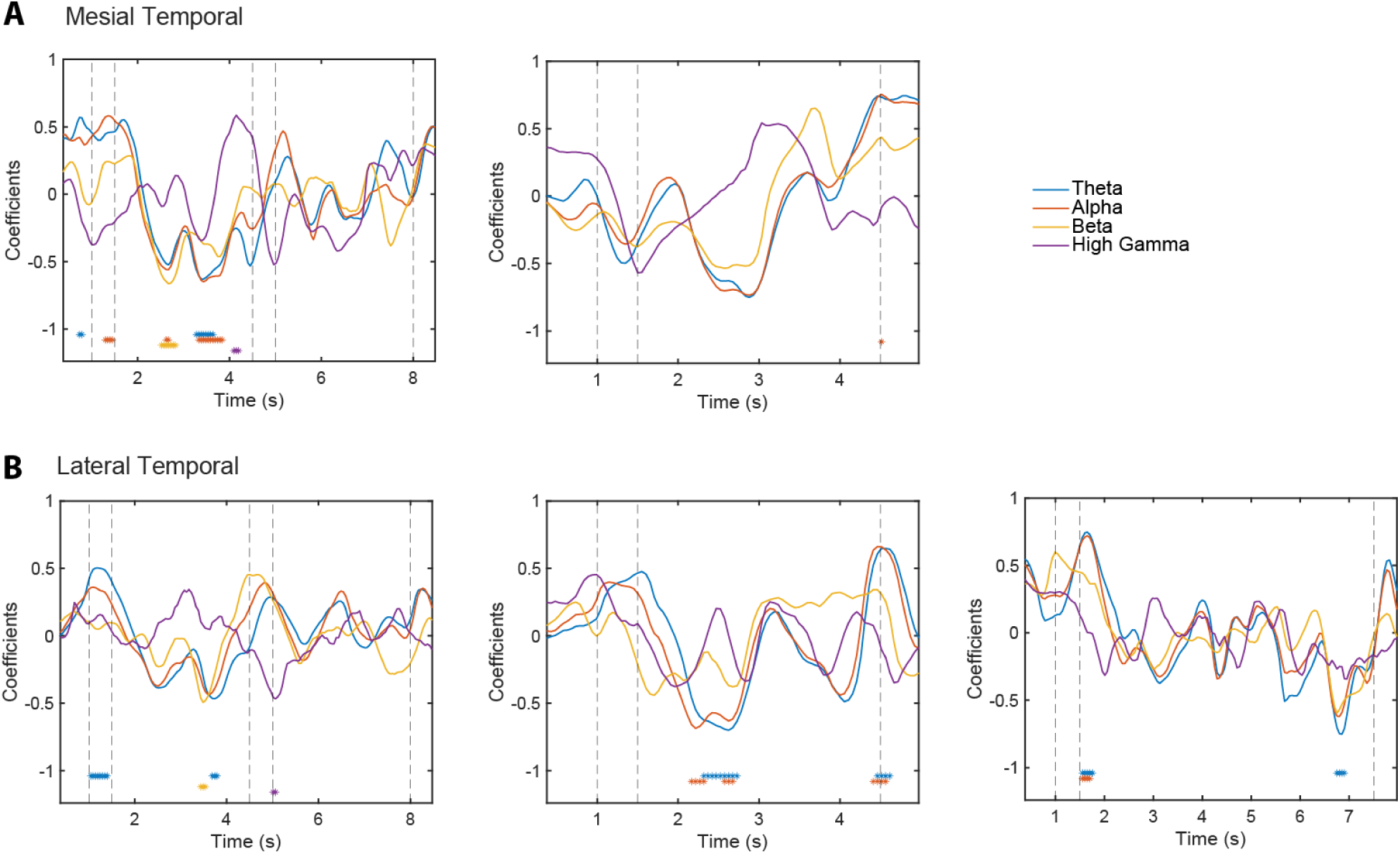
Correlation between neural activity in laboratory tasks and performance in the digit span task in two other brain regions. Data are shown for analyses with more than 8 subjects. A. Mesial temporal cortex (n = 13 for correlation of performance with activity with activity during the shape task; n = 9 for 3s spatial task). B. Lateral temporal cortex (n = 19 for correlation of performance with activity during the shape task; n = 11 for 3s spatial task; n = 11 for 6s spatial task). Conventions are the same as in Fig. 4.

**Supplementary Figure S5.**
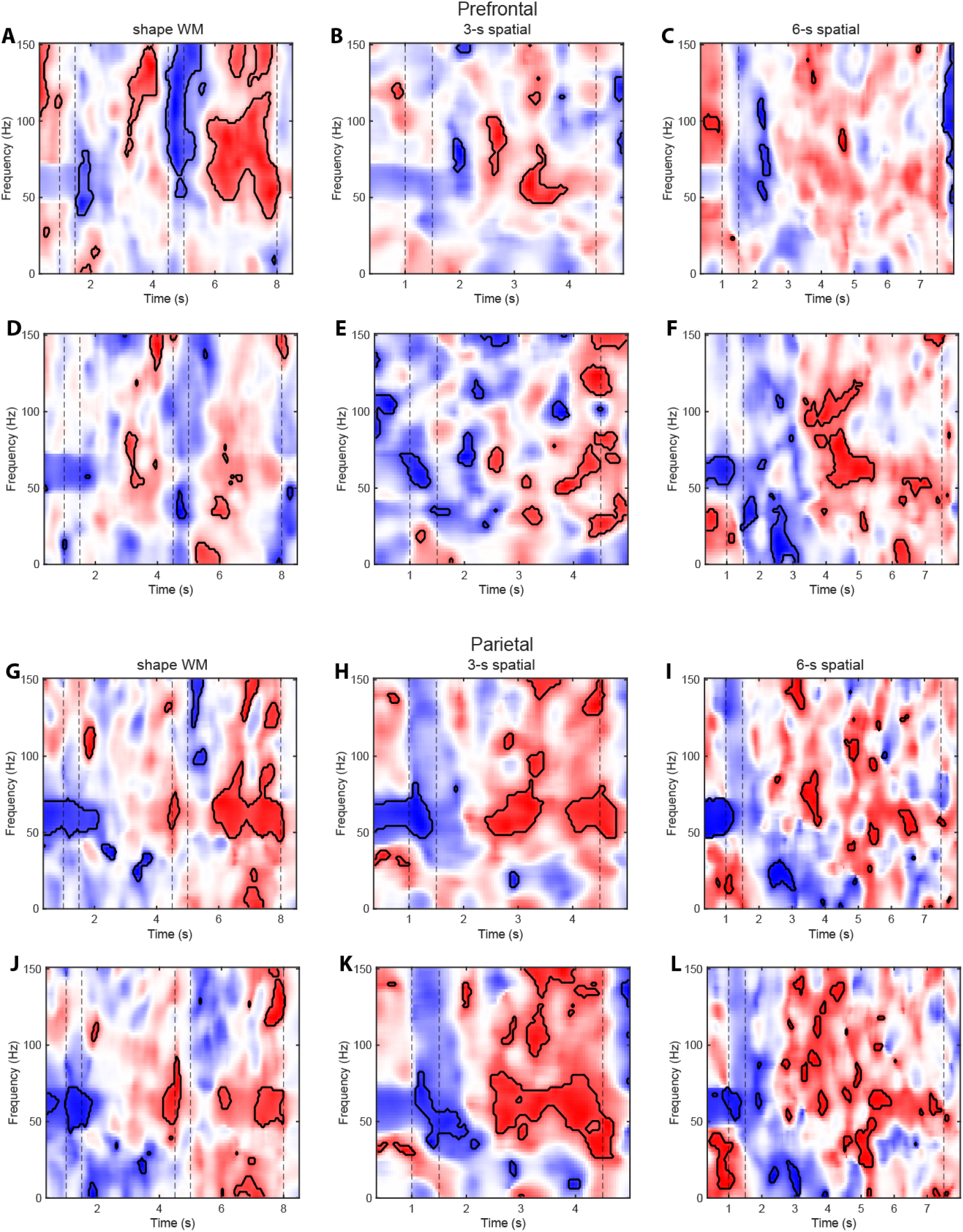
Time-frequency power correlation of prefrontal activity to digit span and mental arithmetic. A-C Time-frequency power correlation of prefrontal activity to digit span performance. D-F. Correlation of prefrontal activity to mental arithmetic performance. G-I. Correlation of parietal activity to digit span performance. J-L. Correlation of parietal activity to mental arithmetic performance. Conventions are the same as in Fig. 5.

**Supplementary Figure S6.**
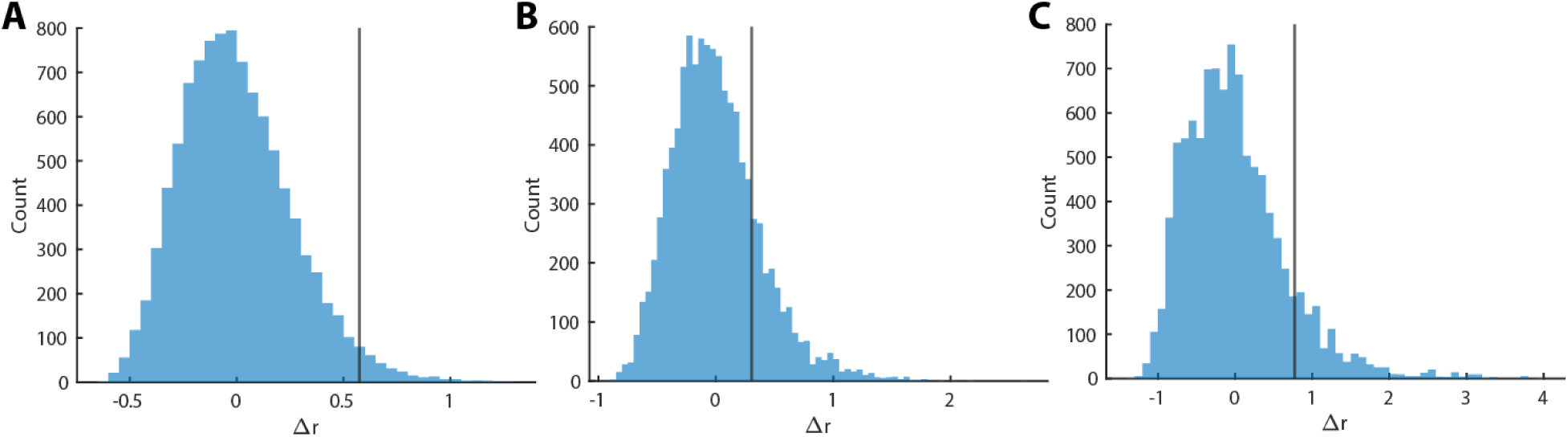
CCA permutation test. Permuted null distribution of the differences between V and U from every time point for the Fisher-transformed correlation coefficients and the null mean for the shape task (A), 3s spatial task (B) and 6s spatial task (C). Line represents difference between Fisher-transformed observed correlation coefficient and null mean.

